# Seed vernalization and gibberellic acid interact to affect life cycle type in facultative winter annual Canadian horseweed (*Erigeron canadensis*)

**DOI:** 10.1101/2025.10.06.680458

**Authors:** Robin Waterman, Brooke Catlett, Ishwari Bhatt, Georgia Edmonds, Jeffrey K. Conner

## Abstract

**Background and Aims:** Plants display enormous variation in the phenological traits that make up their life cycle both within and between populations. Facultative winter annual species are particularly interesting because they can adopt either a fall-emerging/spring-flowering or spring-emerging/summer-flowering life cycle at the population level via evolution or at the individual level via within-generation and transgenerational plasticity. Responses of phenological traits to the environment have often been found to be mediated by changes in hormone levels, especially the growth hormone gibberellic acid (GA).

**Methods:** We conducted growth chamber and greenhouse experiments using the facultative winter annual agricultural weed *Erigeron canadensis* (horseweed) to investigate the interactive effects of genetic variation; parent plant life cycle; and plastic responses to temperature, light, and GA treatments.

**Key Results:** We found that contrary to a prior report, exposing imbibed seeds to 3-4 weeks of cold (i.e., seed vernalization) does not always result in summer annual type growth, with considerable variation found among field-collected seeds from 10 populations. Further, we found that seed vernalization and exogenous application of GA both tended to increase summer annual characteristics, interacting in ways that were largely consistent with the hypothesis that GA is a mechanism for cold-induced life cycle differentiation. Light treatment did not significantly affect life cycle traits, while parent life cycle type had marginal effects on offspring life cycle type. Finally, genetic variation among and within sites explained far less of the variation in life cycle traits than the plastic responses to seed vernalization and GA treatments.

**Conclusions:** Our study proposes that the seasonality of this harmful agricultural weed is influenced by a GA-mediated response to vernalization of seeds during winter, yet highlights the need for further study, given the variability in this response.

## Introduction

Phenology is the timing of recurring events in organismal life cycles. Phenological traits in plants include time of emergence, flowering, and senescence. The practical applications of studying plant phenology have long been recognized by agricultural scientists (Huberman 1941; Chmielewski 2013), and a growing number of studies in recent decades have focused on documenting phenological shifts in response to anthropogenic climate change (Forrest and Miller-Rushing 2010). Such shifts often serve as crucial mechanisms by which plants become or remain well-adapted to their environments. Plants may plastically adjust their phenology in response to environmental cues received during their own (within-generation plasticity) or their parent’s (transgenerational plasticity) lifetime (Auge *et al*. 2017). In addition, plant populations may evolve in response to selection on phenological traits (e.g. Hall and Willis 2006; Donohue *et al*. 2010).

While incremental changes in plant phenology have been widely studied, categorical changes in life cycle type have received less attention. Plants are typically classified as annuals if they complete their life cycle in one year, biennials if they complete their life cycle in two years, or perennials if they complete their life cycle in more than two years. Annual plants can be further subdivided into summer annuals, which emerge in spring and reproduce by fall; and winter annuals, which emerge in fall, overwinter as a rosette, and reproduce from spring to summer. Facultative winter annuals can adopt either a winter or summer annual life cycle in response to different environmental conditions at the level of the population via evolution (Meyer *et al*. 2004; Bloomer and Dean 2017; Charbonneau *et al*. 2018) and at the level of the individual via plasticity (Sans and Masalles 1994; Lu *et al*. 2014). Both mechanisms often co-occur (Best and Mc Intyre 1976; Landers 1995; Garrison *et al*. 2024). Furthermore, nongenetic inheritance (i.e., parental effects) may also play a role in determining offspring life cycle type (Mennan and Ngouajio 2006; Lu *et al*. 2016; Kanomanyanga *et al*. 2025). The effects of all of these factors on life cycle differentiation are rarely studied together, making it difficult to compare their importance or look for interactions between genes and environment.

A promising proximal mechanism by which environmental or genetic variation influences life cycle differentiation is through changes in levels of the plant growth hormone gibberellic acid (GA). This hormone has been found to play a key role in germination, stem elongation, and flowering initiation (reviewed in Gupta and Chakrabarty 2013; and in Shah *et al*. 2023), which are all important parts of life cycle differentiation. Exogenous GA application has been found to override normal photoperiod or temperature requirements for flowering in at least 42 species of rosette-forming summer annual, winter annual, biennial, and perennial plants (reviewed in Lang and Reinhard 1961; and in Zeevaart 1983). One study showed that the exogenous application of GA accelerated the flowering of winter annual, but not summer annual ecotypes of *Raphanus raphanistrum*, suggesting GA upregulation as a mechanism of evolved rapid summer annual flowering (Garrison 2022). Further, *Chrysanthemum morifolium* summer-flowering mutants were found to have higher endogenous levels of GA than normal fall-flowering plants (Dong *et al*. 2017).

In most previously studied facultative winter annual species, dormancy cycling plays an important role in shaping plant life cycles (Baskin *et al*. 1993; Footitt *et al*. 2013; Burghardt *et al*. 2015). In these systems, the timing of seed maturation and/or dispersal influences the dormancy state of seeds and thus their ability to germinate in different seasons. Other facultative winter annual species are not known to exhibit dormancy, and the mechanisms shaping life cycle variation in these species are less clear. One prior study investigated this question using *Erigeron canadensis* (horseweed), subjecting seeds to various environmental conditions in the growth chamber. Their results suggested that winter vs. summer annual life cycle differentiation can be explained as a plastic response to seed vernalization (Schramski *et al*. 2021), where imbibed seeds are exposed to cold conditions that simulate winter, similar to seed stratification but used to accelerate flowering rather than break dormancy (Michaels and Amasino 2000). Despite this compelling finding, it is limited by having few populations, few individuals per population, and only one parental environment, meaning that possible contributions of genetic and parental effects to the response remain unclear.

The present study builds on prior work in *E. canadensis,* using greenhouse and growth chamber experiments to test We made the following hypotheses and predictions: 1) We hypothesized that life cycle type differentiation in horseweed is primarily plastic based on Schramski et al.’s consistent results among their two collection sites, and thus we expected to find little variation explained among or within populations. 2) We hypothesized that plastic growth type differentiation could be explained by a combination of a) parent life cycle type, b) offspring seed temperature, and c) offspring light environment. Specifically, we predicted that summer annual characteristics would be greatest in offspring of winter annual parents, seeds exposed to winter-mimicking seed conditions, and plants grown in brighter conditions given that summer annual plants germinate in the spring with little competitor biomass present and less daily sunlight than in fall (27 - 40 mol·m^−2^·d^−1^ photosynthetic light Mar. to May vs. 12 – 31 mol·m^−2^·d^−1^ Sep. to Nov.; Faust and Logan 2018). 3) We hypothesized that the effects of seed vernalization are at least partially mediated by increased levels of GA. Based on this, we first predicted that plants treated with GA would be more likely to show summer annual characteristics. We then predicted that exogenous GA would have little effect on seed-vernalized plants if vernalization increases endogenous GA levels, and conversely that seed vernalization would have little effect on plants receiving exogenous GA. Overall, our study aims to provide a more complete picture of the factors that influence variation in horseweed life cycle type, serving as a useful case study for future studies of life cycle types in other species and providing insight for managers of this harmful agricultural weed.

## Materials and Methods

### Study System

*Erigeron canadensis* L. (syn. *Conyza canadensis*, Canadian horseweed, or marestail; hereafter horseweed) is a weedy annual plant in the Asteraceae family that is native to Central and North America (Weaver 2001). It now commonly infests agricultural fields and disturbed habitats in temperate zones across the world. Horseweed forms wind-dispersed, primarily self-fertilized seeds (estimated at 96%, Smisek 1995) that have been classified as nondormant or weakly dormant at most (Baskin and Baskin 1988; Buhler and Owen 1997; Karlsson and Milberg 2007). It is considered a facultative winter annual because it can adopt a winter annual or summer annual life cycle. While there seem to be peaks of emergence in the fall and spring, the species has wide emergence and flowering windows (Main *et al*. 2006; Tozzi *et al*. 2014). Plants classified as winter annuals generally form a low-to-the-ground rosette for overwintering, whereas those classified as summer annuals skip the rosette stage by immediately growing upright. However, there is continuous variation in the degree to which plants form a rosette and the life cycle types are not morphologically distinguishable at flowering, since any rosette leaves have senesced by this point. Highly contrasting proportional spring vs. fall emergence has been reported from nearby sites (Main *et al*. 2006), and there does not appear to be a clear latitudinal pattern in life cycle proportions (e.g. 4-24% summer annuals at 36°N, Main *et al*. 2006; 92-100% at 39°N, Davis and Johnson 2008; 38-32% at 45°N, Buhler and Owen 1997). However, these studies demonstrated that the two life cycle types usually co-occur in the same field. There are also observational reports that the summer annual life cycle is becoming more common in Michigan (Schramski *et al*. 2021).

### 2021 field collections and 2022 greenhouse common garden

We conducted this experiment to look for genetic differentiation among populations in life cycle traits in common environmental conditions (Fig. 1A). In the fall of 2021, seeds were collected from 30 individuals from each of 10 sites (except for 1 site with just 23), spread across Michigan’s lower peninsula (see Fig 1A and Table S1 for site details). These 293 field parents were presumed summer annuals given their seed production in fall. The self-fertilized seeds produced by a given parent plant are likely highly homozygous so we refer to them as lines. Seeds were stored in envelopes with desiccant at 5°C until used. In May 2022, approximately 50 seeds per line were sown in 200-cell plug trays and vernalized in a growth chamber set to 4°C and 8-hour daylength for 3 weeks (based on Schramski *et al*. 2021). Trays were then moved to a greenhouse at the Kellogg Biological Station (Hickory Corners, MI), thinned to up to five seedlings per line, and transplanted at the first true leaf stage into 16.5-cm-diameter pots filled with SureMix potting soil (Michigan Grower Products, Inc.). Pots were thinned to one plant when leaves started to substantially overlap. Plants that had not bolted by the end of August (about 11 weeks since movement to greenhouse) were vernalized for 35 days in a growth chamber in the same conditions as those used for seed vernalization. A fungicide soak was applied to the vernalized plants to prevent the spread of fungal root rot. Plants were assigned to one of three categorical growth types based on visual assessment by the same observer about 1 week after emergence and updated about 9 weeks after emergence (as in Schramski et al. 2021). “Rosette” types had a basal circular arrangement of dark green, round leaves; “Upright” types had lighter and more elongated leaves arranged along a visible stem (i.e., caulescent); “Intermediate” types fell somewhere between the Rosette and Upright types in their characteristics. Of the 293 sown lines, 228 produced a plant that survived to flower. We also tested initial dormancy levels of seeds from a random subset of 30 of these greenhouse-grown parents with three seeds per parent (total N = 90) by sowing them in petri dishes, monitoring germination for 2 weeks, then using a tetrazolium dye assay to assess viability of ungerminated seeds.

**Figure 1.**
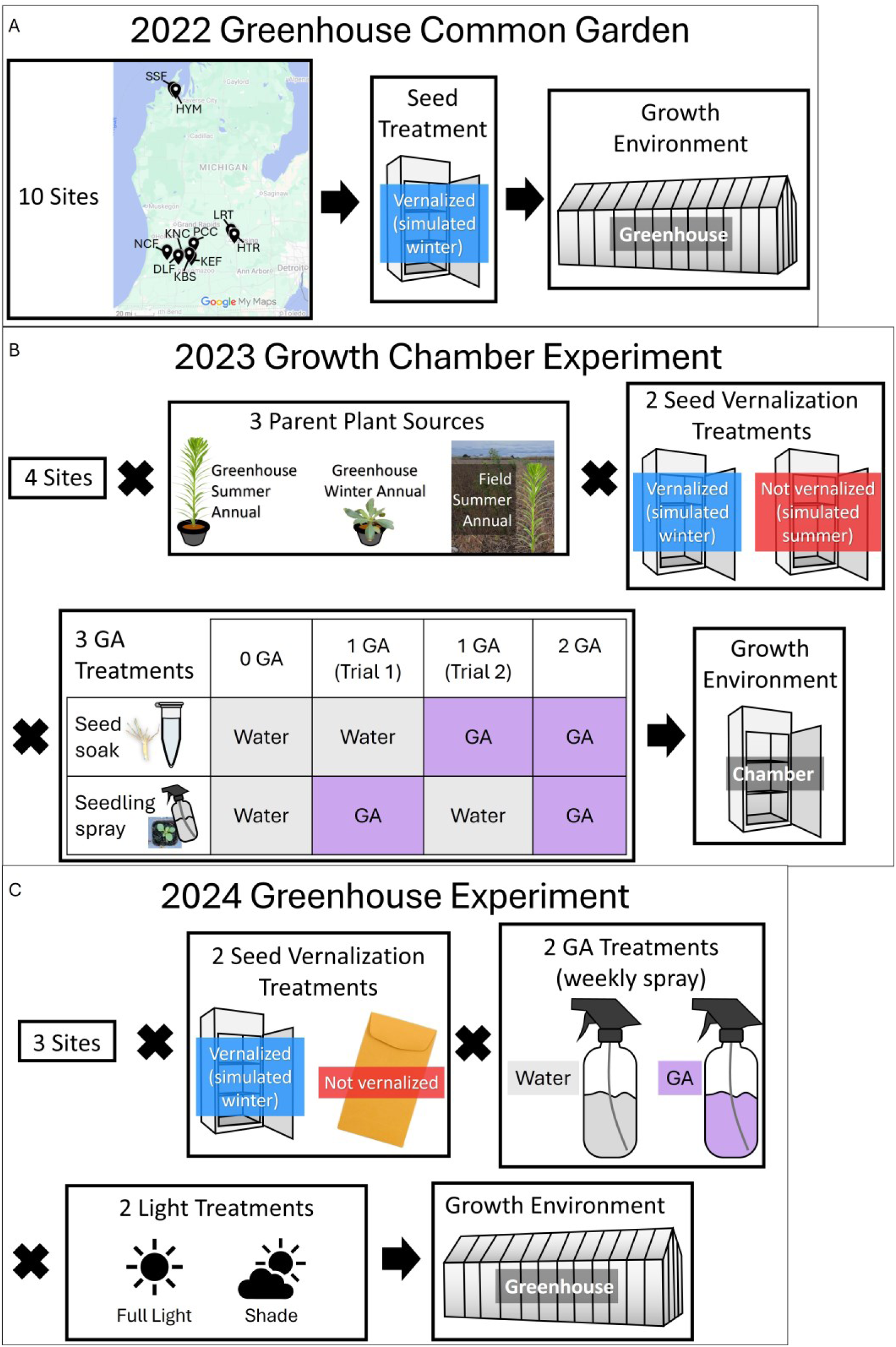
Overview of experimental designs for 2022 Greenhouse Common Garden (A), 2023 Growth Chamber Experiment (B) and 2024 Greenhouse Experiment (C).

### 2023 Growth Chamber Experiment

In this experiment, we tested the effects of seed vernalization and gibberellic acid treatments, along with the seed source factors of population, line, and parent type (field summer annual, greenhouse summer annual, or greenhouse winter annual) on juvenile plant life cycle type traits (Fig. 1B). Seeds were sourced from six lines from each of a subset of four of the sites (DLF, KNC, NCF, and PCC; Table S1). Because we did not have seeds from the same line exhibiting different life cycle types in the same generation, we included offspring of three parent plant groups: 2021 field summer annual, 2022 greenhouse summer annual, and 2022 greenhouse winter annual (those requiring rosette vernalization to flower), with greenhouse-produced seeds briefly stored at room temperature prior to use. Comparing the latter two groups tests for parental effects of life cycle type while controlling for other possible parental effects, but not for effects of genetic variation due to line identity. Comparing the first and last group tests for parental effects of life cycle type while controlling for genetic variation, but not for possible parental effects from the field vs. greenhouse environment. Eighteen replicate seeds from each of the four sites, three families, and three parent source types were randomly assigned to one of two vernalization treatments and one of three GA treatments (detailed below)in two trials, initiated 15 days apart (4 sites x 3 parent types x 3 lines x 2 vernalization treatments x 3 GA treatments x 3 individuals x 2 trials = 1296 total). A total of 846 seeds emerged and survived until data collection (65%).

We tested the effects of GA applied as a seed soak, a cotyledon spray, neither, or both. Both trials included 0, 1 and 2 applications of GA, but due to limited space, we could not include all four combinations of seed soak and leaf spray in both trials. In Trial 1, the single GA application was the seed soak, while in Trial 2 it was the leaf spray. Our results focus on comparing plants that received no GA (“**Water**”) with those that received both the GA seed soak and GA leaf spray (“**GA**”), which was replicated across trials. Seeds were soaked in either a GA solution (1 g/L, Purdom and Glover 2017) or in distilled water for 24 hours at room temperature. At the point of cotyledon emergence, seedling leaves were sprayed with approximately 0.25 mL of either a GA solution (0.035 g/L = 10^-4^ M, Khan et al. 2006) or distilled water.

After completing their soaking treatments, seeds were sown onto the surface of 200-cell plug trays filled with moistened SureMix potting soil, with one tray per treatment (3 GA treatments x 2 vernalization treatments x 2 Trials = 12 total trays) and randomized positions within trays. **Vernalized** seeds were placed in a simulated winter growth chamber set to 4°C and 9-hour daylength. **Unvernalized** seeds were placed in a simulated summer chamber set to 24°C and 15-hour daylength. Although in the field we expect seeds to primarily emerge in fall or spring, the unvernalized seeds sown in simulated summer conditions readily emerged (Fig. S1). Surprisingly, some seeds in simulated winter conditions emerged in the cold (Fig. S1). To prevent cold-related mortality, the vernalization period was ended after about 40% of seeds had emerged, after 3 weeks in Trial 1 and 1 week in Trial 2. After their vernalization treatments, both growth chambers were set to 17°C and 12-hour daylength to simulate spring/fall equinox conditions (based on local average max temperatures and daylengths in mid-fall and mid-spring).

Pots were censused every day for newly emerging seedlings (final seedling emerged on day 33 in Trial 1 and day 18 in Trial 2). Seedlings were defined as having emerged once a stem and both cotyledons were visible. After 3 – 5 weeks of growth, each individual was assigned a categorical growth type (as in 2022 greenhouse common garden). Given the relatively continuous range of variation we observed, we complemented the categorical data with quantitative measures of leaf roundness and plant height growth rate, since rosette-forming winter annuals make rounder leaves and remain close to the soil surface prior to bolting (cite or pers. obs.). Measurements were taken of plant height, plant diameter, and leaf roundness on the same day by vernalization treatment. Vernalized plants were measured 16 days after unvernalized plants, enabling measurements to be taken after the same mean number of days since emergence in Trial 1, but 11 days apart in Trial 2 (staggering measurement points could not be done due to logistical constraints). Height was measured with a ruler as the vertical distance between the soil and the last node (meaning that height was close to 0 for a rosette structure). Overhead photographs and manual tracing in ImageJ were used to measure maximum plant diameter and leaf roundness (4 x area/[pi x major axis^2^], ranging from 0 to 1, with 1 being a perfect circle) on one leaf/plant for all plants with at least one clearly visible entire leaf (N = 785). Height growth rate was calculated as height in mm/days since emergence (results from alternative calculation as height in mm/diameter in mm provided in supplemental). Although some plants were measured on different days since emergence, our analyses examine height growth rate rather than absolute height, and leaf shape rather than size as quantitative indicators of life cycle type.

### 2024 Greenhouse Experiment

In this experiment, we tested the effects of seed vernalization, gibberellic acid, and light treatments, along with the seed source factors of population and line on life cycle traits in plants grown to maturity (Fig. 1C). Seeds were sourced from two to three lines from each of two sites from the 2022 greenhouse generation (HTR and LRT) plus five lines from fall 2023 collections from one of the same fields used in Schramski et al. (2021, MSU Agronomy Farm, Lansing, MI) using the coordinates they provided (Table S1). For each selfed line, pots (7-cm-diameter) were randomly assigned to one of eight treatment combinations, detailed below, with 5 replicate seeds per pot. Pots were filled with SureMix potting soil. Of the 400 seeds planted, 26% emerged and of those that emerged, 46% survived to flower (substantial mortality resulted from fungal root rot).

For **Vernalized** pots, soil was soaked and then the 5 seeds were spread on the surface of the soil. The pots were placed in randomized positions in a growth chamber set to 4°C and 8-hour daylength. Pots were checked weekly for any emerging seedlings and to ensure moist soil conditions. After four weeks, pots were removed from the growth chamber and placed in a greenhouse (Kellogg Biological Station, Hickory Corners, MI). On the same day, **Unvernalized** seeds were planted in pots in the same manner as the vernalized pots and placed in the greenhouse. Note that in this experiment, the vernalization control was untreated seeds because we wanted plants in both treatments to germinate and develop at the same time. Pots were placed on the same greenhouse bench in randomized positions within one of two light treatments: **Shade** pots were placed under a shade tent (shade cloth over PVC pipes, mimicking fall emergence among abundant vegetation), **Full Light** plants were placed in the open (mimicking spring emergence among sparse vegetation). Shaded plants received about 76% less photosynthetically active radiation (based on 20 measurements per treatment with an Accupar LP-80 ceptometer). Plant leaves were sprayed weekly from emergence to flowering with either **Water** (distilled) or **GA** solution (0.035 g GA_3_/L distilled water). All pots were kept well-watered and maintained until all plants had either died or flowered.

Pots were censused every 4 days for newly emerging seedlings for the first 16 days, and every 7 days for the next 7 weeks, at which point no further emergence occurred. Plants were also censused weekly for signs of flowering, and plants were marked as having flowered if they contained at least one fully expanded flower. Plant height was measured as the length of the stem from the base of the soil to the last node on days 21, 49, and 64 since planting, along with at the first census in which they flowered. We defined height growth rate as day 64 plant height in mm/days since emergence. Nine weeks after planting, plants were assigned a categorical growth type (as in prior experiments) and leaf roundness was estimated from photos of the largest leaf from one plant per line.

### Statistical Analyses

All statistical analyses were done in RStudio running R v. 4.4.2 (R Core Team 2024). The package “emmeans” (Lenth 2024) was used to test significance of model terms (Type III Analyses of Deviance), conduct post-hoc analyses on treatment combination means, and generate estimated marginal means. All plots were made using the package “ggplot2” (Wickham 2016). The package “MuMIn” (Bartoń 2025) was used to calculate marginal and conditional R^2^ values for our linear mixed effects models, where marginal is the variance explained by the fixed effects and conditional is the variance explained by both fixed and random effects (Nakagawa and Schielzeth 2013). The difference between conditional and marginal was used to estimate the additional variance in the response explained by adding the random effect of line.

For the 2023 growth chamber experiment, we used linear mixed effects models with the response variables of height growth rate (log-transformed to improve normality of residuals) and leaf roundness; the fixed effects of GA treatment (GA vs. Water), seed vernalization treatment (vernalized vs. unvernalized), trial (1 vs. 2), and all two- and three-way interactions plus parent plant type (greenhouse winter annual vs. greenhouse summer annual vs. field summer annual), seed source site, and the temperature at which a plant emerged (4 vs. 17 vs 24 °C); and the random effect of selfed line. We ran similar models except that we replaced GA treatment with number of GA applications to look for different effects of the GA seed soak vs. leaf spray.

We used a similar approach to model height growth rate, leaf roundness, and days to first flower in the 2024 greenhouse experiment, except that a log transformation was not needed to get normally-distributed residuals, greenhouse light treatment (full light vs. shade) replaced trial, and parent type and emergence temperature were removed. Additionally, for the leaf roundness model we removed the three-way interaction since the model did not converge when it was included, given the reduced sample size from subsampling leaf measurements.

For our categorical growth type variable, given that the frequency of Intermediate was small and unbalanced, we analyzed two binary growth type variables, either grouping Intermediate with Upright or Rosette. Results for the two grouping methods were quantitatively similar so we only report the latter grouping. Logistic regressions did not converge due to probabilities of 1 or 0 in some groups, so instead we ran pairwise two-sample Fisher’s exact tests separately by the fixed effects in our models listed above. For fixed effects with more than two levels, we adjusted *P*-values for multiple testing using the Benjamini-Hochberg procedure to control the False Discovery Rate.

## Results

### Hypothesis 1: Genetic differentiation in life cycle type

In our 2022 greenhouse common garden, the life cycle type of plants sourced from field-collected, vernalized seeds varied significantly among 10 collection sites in Michigan, explaining 15% of the deviance in the response from a saturated model (Fig. S2A, *P* = 4.8E^-5^). However, the patterns do not appear to track latitude or average winter temperatures (Table S1; e.g., lack of similarity between nearby DLF and KNC sites). Because we grew field-collected seeds in a common environment, these differences reflect some combination of evolved genetic differentiation and parental effects.

We then grew offspring of these 2022 greenhouse plants alongside offspring of field parents from a subset of four sites in our 2023 growth chamber experiment. In our 2024 greenhouse experiment we grew greenhouse offspring from a subset of two sites, plus offspring of field parents from a field site used by Schramski et al (2021). In both experiments, source site did not significantly affect any of our measured variables in either experiment (Tables 1 and 2). This lack of a significant site effect held when including only offspring of greenhouse parents (*P* > 0.23).

**Table 1.** Results of Fisher’s exact tests comparing the proportion of plants assigned to the Upright type vs. Intermediate + Rosette types. For variables with more than two levels, the P-value_Adj_ column shows P-values adjusted for multiple testing using FDR.

| Variable | 2023 Growth Chamber Experiment |  |  |  | 2024 Greenhouse Experiment |  |  |  |
| --- | --- | --- | --- | --- | --- | --- | --- | --- |
|  | Comparison | Odds-Ratio | P-value | P-value<sub>Adj</sub> | Comparison | Odds-Ratio | P-value | P-value<sub>Adj</sub> |
| GA | Water vs. GA | 0.000 | 0.000 |  | Water vs. GA | Inf | 0.000 |  |
| Vernalization | Unvernalized vs. Vernalized | 1.000 | 1.000 |  | Unvernalized vs. Vernalized | 5.264 | 0.033 |  |
| Trial | Trial 1 vs. Trial 2 | 1.290 | 0.164 |  |  |  |  |  |
| Parent Type | GH WA vs. Field SA | 0.909 | 0.634 | 0.922 |  |  |  |  |
|  | GH WA vs. GH SA | 0.886 | 0.505 | 0.922 |  |  |  |  |
|  | Field SA vs. GH SA | 0.975 | 0.922 | 0.922 |  |  |  |  |
| Emergence Temperature | 4 vs . 17 | 1.014 | 1.000 | 1.000 |  |  |  |  |
|  | 4 vs. 24 | 1.239 | 0.246 | 0.428 |  |  |  |  |
|  | 17 vs. 24 | 1.221 | 0.285 | 0.428 |  |  |  |  |
| Site | DLF vs. KNC | 0.745 | 0.187 | 0.375 | HTR vs LRT | 3.381 | 0.135 | 0.202 |
|  | DLF vs. NCF | 1.031 | 0.917 | 0.917 | HTR vs MAF | 3.381 | 0.135 | 0.202 |
|  | DLF vs. PCC | 1.122 | 0.621 | 0.832 | LRT vs MAF | 1.000 | 1.000 | 1.000 |
|  | KNC vs. NCF | 1.383 | 0.153 | 0.375 |  |  |  |  |
|  | KNC vs. PCC | 1.507 | 0.062 | 0.370 |  |  |  |  |
|  | NCF vs. PCC | 1.089 | 0.693 | 0.832 |  |  |  |  |
| Light |  |  |  |  | Shade vs. Light | 0.776 | 0.769 |  |

**Table 2.** Model results for quantitative measures of life cycle type. Test statistics are F-ratios for fixed effects and Likelihood-ratios for random effect (Line). R^2^_m_ is R_2_ for only fixed effects, R^2^_c_ is R_2_ for full model.

| Model Term | DF (Num) | DF (Den) | Test Statistic | P-value | Model Term | DF (Num) | DF (Den) | Test Statistic | P-value | Model Term | DF (Num) | DF (Den) | Test Statistic | P-value |
| --- | --- | --- | --- | --- | --- | --- | --- | --- | --- | --- | --- | --- | --- | --- |
| 2023 Growth Chamber Experiment |  |  |  |  |  |  |  |  |  |  |  |  |  |  |
| **Height Growth Rate** | $R^2_m =$ | 0.504 | $R^2_c =$ | 0.520 | **Leaf Roundness** | $R^2_m =$ | 0.475 | $R^2_c =$ | 0.498 | | | | | |
| GA | 1 | 536.9 | 289.4 | 0.0000 | GA | 1 | 493.0 | 293.6 | 0.0000 |  |  |  |  |  |
| Trial | 1 | 538.6 | 6.3 | 0.0123 | Trial | 1 | 496.7 | 96.8 | 0.0000 |  |  |  |  |  |
| Vernalization | 1 | 542.2 | 2.0 | 0.1571 | Vernalization | 1 | 498.4 | 3.9 | 0.0489 |  |  |  |  |  |
| Parent Type | 2 | 24.7 | 4.2 | 0.0275 | Parent Type | 2 | 23.8 | 3.6 | 0.0441 |  |  |  |  |  |
| Site | 3 | 16.6 | 1.4 | 0.2876 | Site | 3 | 16.6 | 0.4 | 0.7451 |  |  |  |  |  |
| Emergence Temp | 1 | 543.6 | 48.2 | 0.0000 | Emergence Temp | 1 | 499.4 | 2.3 | 0.1277 |  |  |  |  |  |
| GA x Trial | 1 | 538.6 | 0.4 | 0.5222 | GA x Trial | 1 | 494.6 | 11.5 | 0.0008 |  |  |  |  |  |
| GA x Vernal | 1 | 539.1 | 37.9 | 0.0000 | GA x Vernal | 1 | 494.7 | 25.6 | 0.0000 |  |  |  |  |  |
| Trial x Vernal | 1 | 534.1 | 8.1 | 0.0047 | Trial x Vernal | 1 | 491.5 | 13.2 | 0.0003 |  |  |  |  |  |
| GA x Trial x Vernal | 1 | 538.1 | 0.1 | 0.7788 | GA x Trial x Vernal | 1 | 495.6 | 0.0 | 0.9972 |  |  |  |  |  |
| Line | 1 |  | 2.6 | 0.1090 | Line | 1 |  | 5.3 | 0.0217 |  |  |  |  |  |
| 2024 Greenhouse Experiment |  |  |  |  |  |  |  |  |  |  |  |  |  |  |
| **Height Growth Rate** | $R^2_m =$ | 0.628 | $R^2_c =$ | 0.734 | **Leaf Roundness** | $R^2_m =$ | 0.617 | $R^2_c =$ | 0.666 | **Days to First Flower** | $R^2_m =$ | 0.732 | $R^2_c =$ | 0.800 |
| GA | 1 | 46.6 | 22.4 | 0.0000 | GA | 1 | 17.2 | 7.5 | 0.0138 | GA | 1 | 34.1 | 12.7 | 0.0011 |
| Vernalization | 1 | 46.2 | 4.5 | 0.0390 | Vernalization | 1 | 15.3 | 2.1 | 0.1658 | Vernalization | 1 | 34.3 | 42.7 | 0.0000 |
| Light | 1 | 46.8 | 41.6 | 0.0000 | Light | 1 | 16.6 | 2.6 | 0.1286 | Light | 1 | 35.0 | 0.3 | 0.5761 |
| Site | 2 | 3.4 | 1.4 | 0.3668 | Site | 2 | 3.0 | 0.4 | 0.6842 | Site | 2 | 2.7 | 2.9 | 0.2129 |
| GA x Vernal | 1 | 47.9 | 5.1 | 0.0286 | GA x Vernal | 1 | 17.3 | 0.5 | 0.4910 | GA x Vernal | 1 | 37.2 | 5.2 | 0.0287 |
| GA x Light | 1 | 47.3 | 18.1 | 0.0001 | GA x Light | 1 | 17.8 | 0.7 | 0.4043 | GA x Light | 1 | 35.5 | 0.2 | 0.6485 |
| Vernal x Light | 1 | 46.0 | 0.6 | 0.4580 | Vernal x Light | 1 | 16.5 | 0.0 | 0.9699 | Vernal x Light | 1 | 34.4 | 0.0 | 0.9430 |
| GA x Vernal x Light | 1 | 47.1 | 0.4 | 0.5428 |  |  |  |  |  | GA x Vernal x Light | 1 | 35.2 | 0.6 | 0.4595 |
| Line | 1 |  | 1.2 | 0.2714 | Line | 1 |  | 0.1 | 0.7240 | Line | 1 |  | 1.9 | 0.1724 |

In both experiments, we also included multiple genetic lines from each site, expected to be highly homozygous in this selfing species. Line explained a significant amount of variation in leaf roundness only in the growth chamber experiment. Even so, line explained only 2% of the variance, compared with 37% explained by GA and vernalization treatments for height growth rate and leaf roundness, respectively. The lack of variation in life cycle type explained by variation between and within populations is consistent with the hypothesis that life cycle is primarily determined via plasticity.

### Hypothesis 2a: Plastic response to parent life cycle type

In the 2023 growth chamber experiment we included seeds generated by parents exhibiting the summer annual growth type in the greenhouse, winter annual growth type in the greenhouse, or summer annual growth type in the field. We predicted that offspring of summer annual parents would be more likely to exhibit a winter annual life cycle than offspring of winter annual parents, enabling an alternation of life cycles. Parent type had small but significant effects on height growth rate and leaf roundness (Table 2), where offspring of field summer annuals had slightly greater winter annual characteristics than the other two parent types, but only significantly so for one of two comparisons for each trait (Fig. S3). Categorical assignment to the upright summer annual type was similar among the three parent types (Table 1). Thus, while there was a slight difference between the field and greenhouse parents for our quantitative measures, the patterns do not support an effect of parent life cycle type.

### Hypothesis 2b: Plastic responses to seed vernalization

Contrary to our prediction that all cold-vernalized seeds would develop as summer annuals, when seeds were vernalized for three weeks and grown in a common greenhouse environment in 2022, an average of 65% of plants formed an overwintering rosette structure indicative of a winter annual life cycle. Similarly, 66% of plants only flowered after receiving additional cold through rosette vernalization, after failing to bolt for 11 weeks (87% of these were Rosette types; Fig. S2A).

In the 2023 and 2024 experiments, the overall effects of seed vernalization were also variable, but tended to increase summer annual type characteristics, in line with our prediction. Specifically, seed vernalization increased summer annual characteristics for all measures in the greenhouse experiment (though not significantly in the case of leaf roundness; Fig. 2B, Fig. 3B,D,E). In the growth chamber experiment, 3-week (Trial 1) or 1-week (Trial 2) seed vernalization significantly affected leaf roundness in the predicted direction (Fig. 3C) but did not significantly affect categorical growth type assignment or height growth rate (Fig. 2A and Fig. 3A).

**Figure 2.**
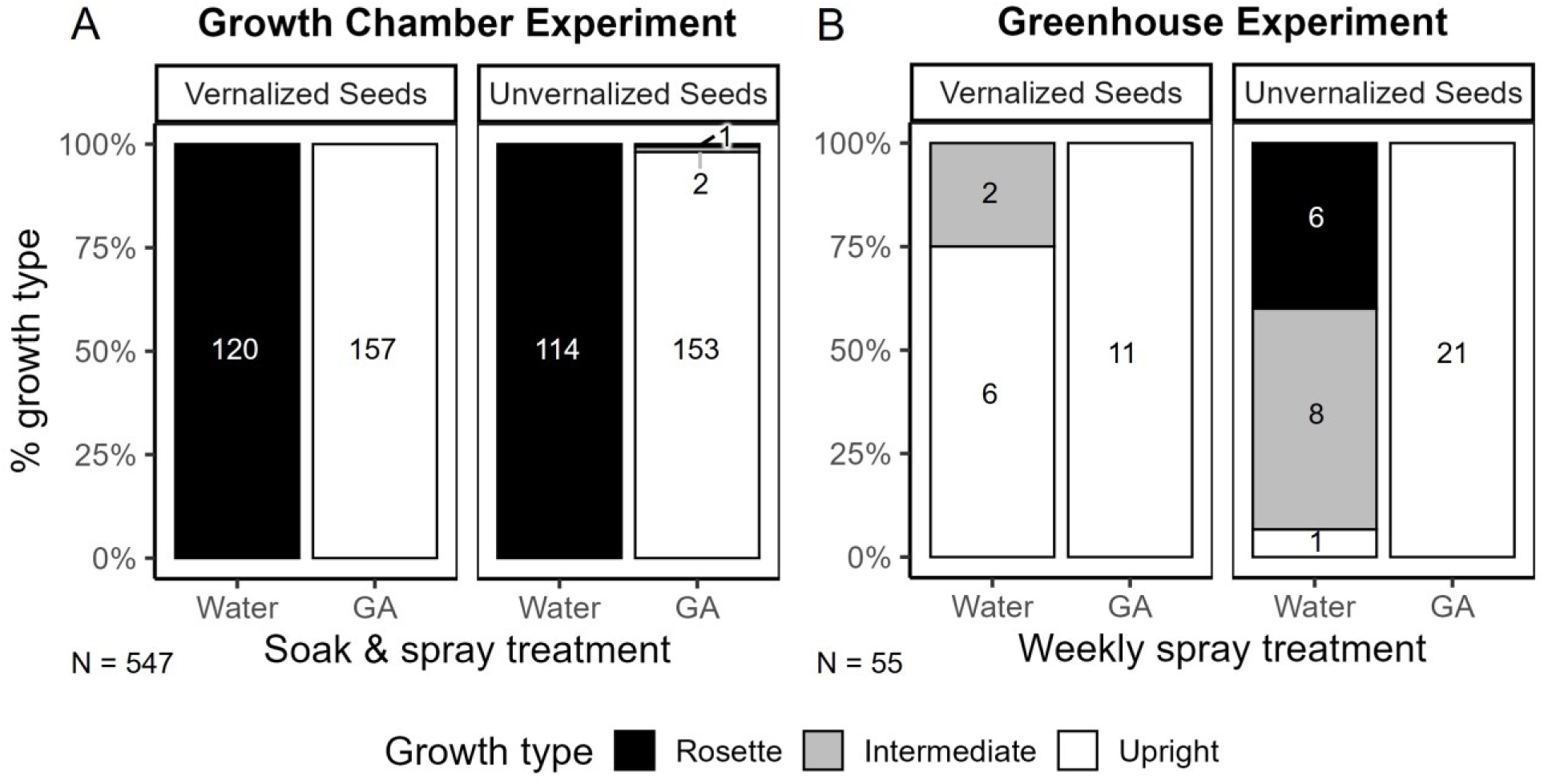
Qualitative measure of life cycle growth type in 2023 Growth Chamber (A) and 2024 Greenhouse (B) Experiments. Colored bars show the proportion of plants categorically assigned to Rosette (black), Intermediate (grey), or Upright (white) growth, with counts shown within bars. Panels separate seed vernalization treatments. Numbers in bars are counts.

**Figure 3.**
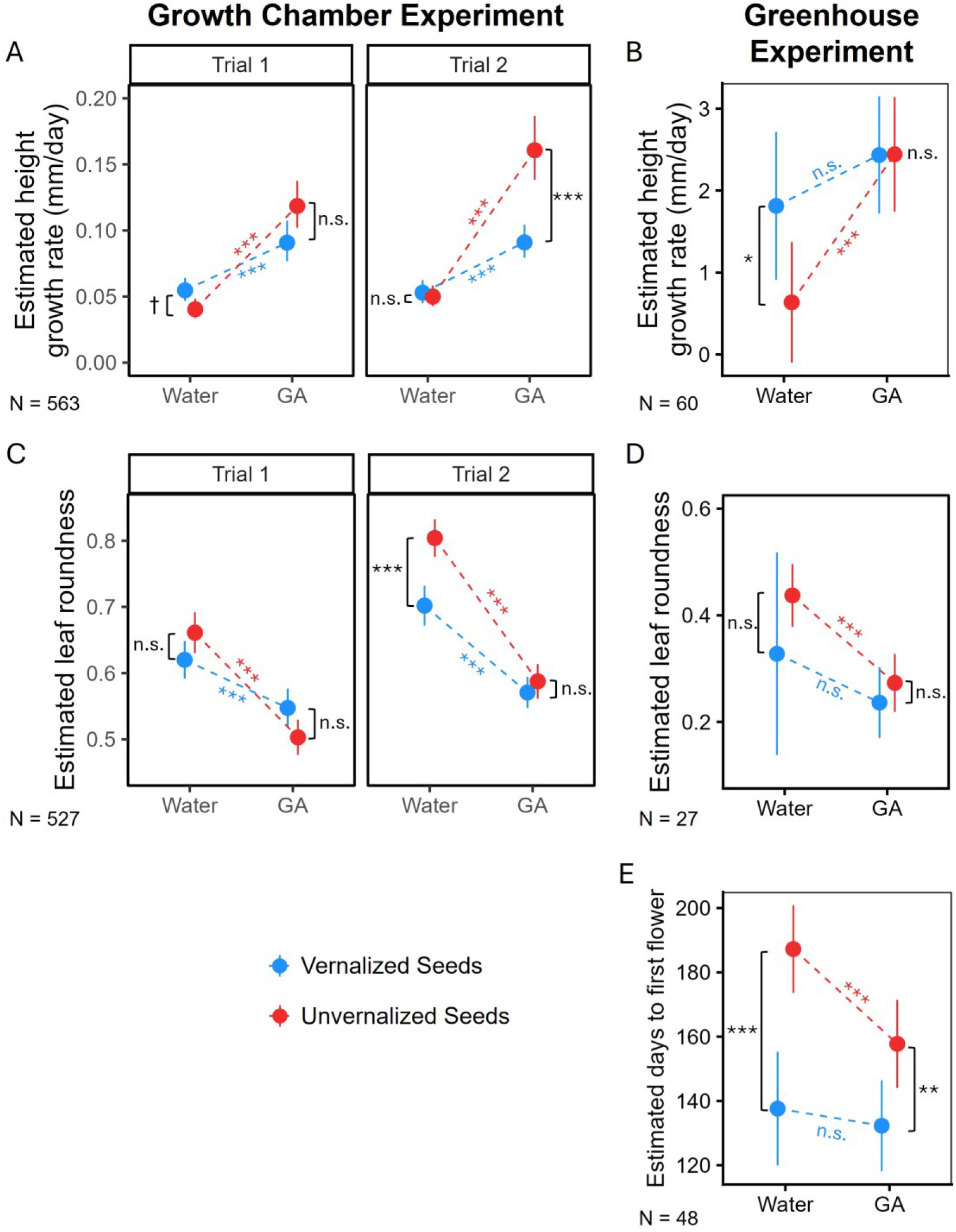
Quantitative measures of life cycle type in 2023 Growth Chamber (A and C) and 2024 Greenhouse (B,D, E) Experiments: height growth rate (A and B), leaf roundness (C and D), and flowering time (E). Colors separate seed vernalization treatments and the slope of dotted lines is the effect of adding GA within vernalization treatment. Panels separate trials in A and C. Points are estimated marginal means after accounting for the other effects in the model and error bars are 95% confidence intervals. Asterisks on lines indicate significance of GA effect within vernalization groups, while asterisks between points indicate significance of vernalization effect within GA treatment groups from Tukey post-hoc tests. ^n.s.^*P* ≥ 0.10, ^†^*P* <0.10, \**P* < 0.05, \*\**P* < 0.01, \*\*\**P* < 0.001.

As expected for seeds lacking dormancy, the cold seed vernalization treatment had little effect on emergence rates, with the effect in opposite directions in the two experiments (+5% and -10% emergence of vernalized seeds in 2023 and 2024, respectively; Fig. S4A and B). Also consistent with the idea that horseweed seeds have little to no dormancy, our seed viability test of a subset of greenhouse-grown seeds showed that 97% of seeds either germinated or were nonviable, while 3% did not germinate and were still viable (Fig. S4D).

### Hypothesis 2c: Plastic response to offspring light environment

In the 2024 greenhouse experiment, we tested our prediction that shaded plants would be more likely to develop winter annual characteristics, particularly if seeds were not vernalized; that is, mimicking late summer or fall germination of seeds produced that year. However, light treatment did not significantly affect growth type assignment, leaf roundness, or days to first flower, and did not significantly interact with vernalization treatment for any measure (Tables 1 and 2). While shaded plants had lower overall height growth rate (-1.7 mm/day) and were less affected by GA spray, given that light treatment did not affect any other measure of life cycle type, we interpret this result as simply reflecting the stunted growth expected from light-limited plants, rather than the limited elongation of rosette-forming plants.

### Hypothesis 3: Seed vernalization response mediated by increased GA

In line with our predictions, the application of GA strongly and significantly increased all of our measures of summer annual type growth across both experiments. Specifically, plants exposed to exogenous GA were more likely to be assigned the Upright growth type (Fig. 2, Table 1), extended their height at a faster rate (Fig. 3A,B), had more elongated leaves (Fig. 3C,D), and flowered sooner (Fig. 3E). This pattern held in the growth chamber experiment when excluding plants in the vernalization treatment that unexpectedly emerged while still in the cold (Fig. S4) or when quantifying growth rate as plant height per plant diameter rather than per day (Fig. S5). Across our life cycle measures, applying GA only to seeds rather than only to leaves generally had a weaker effect in the same direction (Fig. S6), but this comparison was not replicated and so should be interpreted with caution. In line with our expectation for nondormant seeds, the GA seed soak had little effect on emergence rates (water control = 75% vs. GA soak = 79%, Fig. S4C).

We also predicted that if increased GA level is a mechanism by which seed vernalization influences life cycle type, then the two treatments would be at least partially redundant. This would manifest as a statistical interaction between GA and vernalization treatments whereby each factor would have a stronger effect in the absence of the other. We found this predicted pattern for all measures in the greenhouse, but only for leaf roundness in the growth chamber. Specifically, in the greenhouse experiment, seed vernalization in the absence of GA or GA in the absence of seed vernalization caused nearly all plants to be assigned Upright (interaction not statistically testable, Fig. 2B). For leaf roundness in both experiments, along with flowering time and height growth rate in the greenhouse experiment, the vernalization effect was larger in magnitude in the water control group compared with the GA-treated group (compare differences between red and blue points within treatments in Fig. 3B-E; although note that this was not significant for leaf roundness in the greenhouse). Similarly, for these measures, the GA effect was larger in magnitude in the unvernalized group than the vernalized group (compare slopes of red and blue lines in Fig. 3B-E). However, seed vernalization did not affect growth type assignment within either the GA or Water groups in the growth chamber experiment (Fig. 2A). For height growth rate in the growth chamber experiment, the magnitude of the GA effect was larger in unvernalized seeds as predicted, but the vernalization effect switched from slightly positive in the water-treated group (+0.009 mm/day, *P =* 0.07) to negative in the GA-treated group (-0.05 mm/day, *P* < 0.0001), opposing our prediction. Although the three-way interaction was not significant, we note that this surprising negative effect of vernalization on height growth rate in in the GA-treated group was only apparent in Trial 2, where seeds were only vernalized for 1 week (Fig. S1).

## Discussion

### Life cycle type differentiation is primarily plastic rather than genetic

We found that field-collected *E. canadensis* seeds from 10 Michigan sites vernalized and grown in a common greenhouse environment varied between and within sites in morphological growth type and requirement for further vernalization of rosettes. However, when we grew offspring of plants from subsets of these sites plus new collections from a site used by Schramski *et al*. (2021), site differences largely disappeared. For example, sites KNC and DLF were some of the most divergent in the 2022 greenhouse but did not significantly differ in the 2023 growth chamber. Although we only grew seeds from six of the 10 originally sampled sites, our results suggest that the differences in traits associated with life cycle type found in the first greenhouse generation primarily reflect nongenetic parental effects, rather than evolved genetic differentiation. The low levels of differentiation between presumably highly homozygous families further supports the idea that variation in life cycle type is not strongly predicted by genetic variation. Although Schramski *et al*. (2021) did not statistically test for differences among their four seed families from each of four fields, the highly consistent responses of all plants to their experimental treatments suggests little genetic variation for life cycle type, consistent with our study. However, genetic differentiation across a broader geographic range in this widely distributed weed remains possible and should be tested in future studies.

### Mixed evidence for nongenetic effects of parent life cycle type

In addition to comparing field-collected vs. greenhouse-grown seeds, we also explicitly looked for effects of parental life cycle type on offspring life cycle type in our growth chamber experiment. We predicted that differences in the conditions experienced by summer annual versus winter annual parents would reduce parent-offspring resemblance. Alternating parent-offspring life cycles might be adaptive in nature if the parent environment is strongly predictive of the offspring environment (Uller 2008) whereby seeds are shed by winter annual parents in favorable conditions for summer annual emergence, and vice versa. A similar pattern has been found previously in the facultative winter annual *Isatis violascens,* where parent germination season biases seeds towards the dimorphic dormancy type that promotes germination in a different season than the parent (Lu *et al*. 2016). First, our study confirms that parents of both life cycle types can produce offspring of both life cycle types, whereas this was previously only shown for summer annual type parents (Schramski *et al*. 2021). We found evidence for a parental effect in the predicted direction for leaf roundness but not height growth rate. Future studies should include field winter annual parents as a comparison group to provide a more complete picture of how parental effects may influence life cycle type. Overall, these results suggest that nongenetic inheritance plays some role in life cycle type differentiation, but the mechanisms (e.g. transgenerationally plastic response to specific environmental cues, indirect genetic effects from altered parental gene expression) shaping these responses remain unclear.

### Some evidence for a plastic response of life cycle type to seed vernalization but no evidence for a plastic response to light level

Given that summer annuals emerge in spring after cold and wet winter conditions, we expected seeds exposed to simulated winter conditions to emerge as summer annuals. Consistent with this expectation, we found that exposing seeds to cold and wet conditions increased summer annual growth characteristics, but not in all plants. In our greenhouse common garden in which all seeds were vernalized for three weeks, only about a third of plants developed as summer annuals. Among plants that did not receive GA in our two experiments, all plants developed winter annual characteristics regardless of vernalization treatment in the growth chamber, yet seed vernalization increased the chances of developing summer annual characteristics in our smaller sample of greenhouse plants. These results contrast markedly from those of Schramski *et al*. (2021), who classified no unvernalized plants as summer annuals, but 88% and 100% of plants vernalized for two weeks and four weeks as summer annuals, respectively. That study and all seed vernalization trials in the present study utilized the same temperature (4°C) and the same or similar photoperiod (8 – 9 hours). These results demonstrate that while seed vernalization plays a role in life cycle determination, its variable effects point to interactions with other environmental or genetic factors.

One possible interacting environmental factor we tested was the amount of available light. We expected cold temperatures followed by higher light availability to result in an increased likelihood of summer annual growth, given that spring-emerging plants face less competitor biomass and receive more mean daily light than fall-emerging plants. However, results from our greenhouse experiment did not support this prediction. While we only manipulated total light level in our experiment, manipulations of light quality and/or daylength might produce different results. A systematic field survey documenting the occurrence of winter vs. summer annual horseweed across its range, along with habitat characteristics where these plants are found (e.g. mean winter temperatures, land use type and management) could suggest additional environmental factors that play a role in life cycle type differentiation and would provide key baseline data for documenting changes in proportional life cycle type occurrence over time. Studies tracking the outcomes of replicate seeds planted in fields varying in known environmental conditions would also be useful.

Overall, the plastic ability of facultative winter annuals to adjust their life cycle to suit their current seasonal environment may be adaptive. Preliminary data suggest that plants expressing the seasonally mismatched growth type (rosette in summer annual field and upright in winter annual field) have lower survival (N = 23 total flowered/247 transplanted seedlings). A highly flexible life cycle within populations may be particularly beneficial in crop fields that are actively managed to prevent horseweed growth. This is because unpredictable and high-mortality environments are expected to favor strategies that produce temporal phenological variation, such as bet-hedging dormancy in other systems (Donohue *et al*. 2010).

### Gibberellic Acid regulation may be a mechanism for horseweed life cycle type differentiation

Alongside our tests of genetic and environmental factors influencing horseweed life cycle type differentiation, we also sought to test a hypothesized mechanism by which such differentiation may occur. In line with our predictions, exogenous application of the growth hormone Gibberellic Acid (GA) increased summer annual characteristics. Also as predicted, this GA effect was stronger in unvernalized plants, while the effects of seed vernalization (in cases where it also increased summer annual characteristics) were stronger in plants not receiving GA. These results suggest that increased levels of GA may be an important mechanism by which plants plastically respond to experiencing a period of cold prior to germination. Greater GA biosynthesis, more efficient GA transport, and decreased GA degradation all might contribute to increasing experienced levels of GA in plants that experienced cold conditions as a seed. Increased GA could accelerate flowering by triggering an earlier initiation of the reproductive growth stage (including immediate bolting upon emergence) or by increasing overall growth rates. The changes in height extension, leaf shape, and rosette formation we observed in our experiments suggest the first possibility. Although we did not explicitly attempt to quantify overall growth, we observed qualitatively similar leaf counts between rosette and upright plants that emerged on the same day.

It remains unclear why we did not find the predicted effect of seed vernalization and its interaction with GA for height growth rate in the growth chamber experiment, which was one of seven measures of life cycle type across both experiments. One possibility relates to the germination of many seeds during seed vernalization at 4°C, which was surprising given prior studies reporting base germination temperatures of 8-14°C (Steinmaus *et al*. 2000; Tozzi *et al*. 2014) and the fact that it never occurred in the common garden or greenhouse experiment. Although results were similar when excluding plants that emerged in the cold, all or a larger fraction of seeds may have begun the germination process, changing their development. We note that the height growth rates for these plants were very low in comparison with the greenhouse plants (mean = 0.1 vs. 2.0 mm/day), potentially indicating stunted growth.

It has long been hypothesized that GA and vernalization act through a common mechanism, given the ability of exogenous GA to initiate flowering in plants that normally require vernalization (Zeevaart 1983). This mechanism has been studied in detail in the model plant *Arabidopsis thaliana*, where vernalization and GA both act on the floral integrator gene SOC1 (Moon *et al*. 2003). Interactive effects of GA and cold treatments on plant life cycle traits have also been studied in a number of crop systems (Mutasa-Göttgens *et al*. 2010; Rezaee *et al*. 2023; Zhao *et al*. 2023), but seldom in weeds. However, Garrison (2022) found that the native winter annual ecotype of *Raphanus raphanistrum* could be induced to flower without rosette vernalization by exogenous application of GA, whereas the weedy summer annual ecotype showed little response to GA. Syntheses on the interactive effects of temperature and GA do not generally distinguish between a cold period at the seed versus the seedling stage, although Chouard (1960) reported that GA seed treatment cannot replace seed vernalization.

Our study extends prior work by suggesting that horseweed seeds experiencing cold winter conditions increase their levels of GA, helping to stimulate development as a summer annual. Interestingly, applying GA at the seedling rather than at the seed stage was generally more effective in producing plants with summer annual traits. A similar tendency for GA spray compared with seed soak to cause accelerated life cycle traits has been found previously in three annual crops (Chakravarti 1958; Wilson 1981). This indicates that although the key environmental cue of cold temperatures may be experienced as a seed, the seedling stage may be a critical period for hormonal regulation. Further studies in other facultative winter annual species are needed to determine the generality of a GA-mediated seed vernalization mechanism for life cycle differentiation. In addition, future studies might test for increases in endogenous levels of GA or expression levels of genes in the GA response pathway in summer annual vs. winter annual plants growing in the field. Finally, comparing the life cycle characteristics of GA-deficient mutants with control plants subjected to seed vernalization vs. unvernalized control treatments would more directly link GA to a vernalization-mediated plastic life cycle response.

### Applied significance

A review of 19 common Canadian winter annual agricultural weeds found that the facultative ability to adopt a winter or summer annual life cycle predominates in these species, yet the authors noted a general lack of information on many important aspects of their biology (Cici and Van Acker 2009). Gaining a better understanding of the genetic and environmental factors contributing to variation in horseweed life cycle types has implications for the management of this widespread and problematic agricultural weed. According to the most recent surveys of US and Canadian farmers from the Weed Science Society of America, horseweed is among the top five most troublesome weeds in winter cereal grains, fruits, nuts, and soybeans (Van Wychen 2022, 2023). Of particular note for management, summer annual type plants, and especially those of resistant biotypes, were found to be more resistant to glyphosate when grown in a greenhouse (Schramski *et al*. 2021). This increased resistance was partially attributable to reduced glyphosate retention (Fisher *et al*. 2023). Prior studies in horseweed and other facultative winter annual populations have also found that by skipping the overwintering stage, spring-emerging summer annuals are more likely to survive to seeding but produce fewer seeds on average than fall-emerging winter annuals (Regehr and Bazzaz 1979; Marks and Prince 1981; Sans and Masalles 1994). Our study indicates that horseweed plants expressing either life cycle type may plastically switch to the other type in the next generation, potentially completing two generations per year. Therefore, switching crop life cycle or eliminating only those horseweed plants with the same life cycle type as the desired crop are unlikely to prevent horseweed infestation. Manipulating the winter temperatures or levels of GA experienced by horseweed seeds may allow managers to bias life cycle type differentiation in a favorable direction. Recently observed increases in the summer annual type in Michigan are less likely to be due to rapid evolution than to a change in the environment or the differential survival of the two life cycle types.

## Author Contributions

All authors helped design the experiments; RW, BC, GE, and IB conducted experiments; RW analyzed the data; RW and BC drafted the manuscript; all authors reviewed and approved the final manuscript.

## Acknowledgements

We thank Kate Shaw and Mark Hammond for greenhouse and growth chamber management; Ava Garrison for assistance with watering; and Second Spring Farm, Natural Cycles Farm, and DeLano Farms for seed collection access. This manuscript was improved by comments from the Conner Lab and XX anonymous reviewers. This is KBS contribution no. XX.

## Literature Cited

Auge GA, Leverett LD, Edwards BR, Donohue K. 2017. Adjusting phenotypes via within-and across-generational plasticity. New Phytologist 216: 343–349.

Bartoń K. 2025. MuMIn: multi-model inference.

Baskin CC, Baskin JM. 1988. Germination ecophysiology of herbaceous plant species in a temperate region. American Journal of Botany 75: 286–305.

Baskin CC, Chesson PL, Baskin JM. 1993. Annual seed dormancy cycles in two desert winter annuals. The Journal of Ecology 81: 551.

Best K, Mc Intyre G. 1976. Studies on the flowering of *Thlaspi arvense* L. III. The influence of vernalization under natural and controlled conditions. Botanical Gazette 137: 121–127.

Bloomer R, Dean C. 2017. Fine-tuning timing: natural variation informs the mechanistic basis of the switch to flowering in *Arabidopsis thaliana*. Journal of Experimental Botany 68: 5439–5452.

Buhler DD, Owen MD. 1997. Emergence and survival of horseweed (*Conyza canadensis*). Weed Science 45: 98–101.

Burghardt LT, Metcalf CJE, Wilczek AM, Schmitt J, Donohue K. 2015. Modeling the influence of genetic and environmental variation on the expression of plant life cycles across landscapes. The American Naturalist 185: 212–227.

Chakravarti S. 1958. Gibberellic acid and vernalization. Nature 182: 1612–1613.

Charbonneau A, Tack D, Lale A, et al. 2018. Weed evolution: Genetic differentiation among wild, weedy, and crop radish. Evolutionary Applications 11: 1964–1974.

Chmielewski F-M. 2013. Phenology in agriculture and horticulture In: Schwartz MD, ed. Phenology: an integrative environmental science. Dordrecht: Springer Netherlands, 539–561.

Chouard P. 1960. Vernalization and its relations to dormancy. Annual Review of Plant Physiology 11: 191–238.

Cici SZH, Van Acker RC. 2009. A review of the recruitment biology of winter annual weeds in Canada. Canadian Journal of Plant Science 89: 575–589.

Davis VM, Johnson WG. 2008. Glyphosate-resistant horseweed (*Conyza canadensis*) emergence, survival, and fecundity in no-till soybean. Weed Science 56: 231–236.

Dong B, Deng Y, Wang H, et al. 2017. Gibberellic acid signaling is required to induce flowering of chrysanthemums grown under both short and long days. International Journal of Molecular Sciences 18: 1259.

Donohue K, Rubio De Casas R, Burghardt L, Kovach K, Willis CG. 2010. Germination, postgermination adaptation, and species ecological ranges. Annual Review of Ecology, Evolution, and Systematics 41: 293–319.

Faust JE, Logan J. 2018. Daily light integral: a research review and high-resolution maps of the United States. HortScience 53: 1250–1257.

Fisher JL, Sprague CL, Patterson EL, Schramski JA. 2023. Investigations into differential glyphosate sensitivity between two horseweed (*Conyza canadensis*) growth types. Weed Science 71: 22–28.

Footitt S, Huang Z, Clay HA, Mead A, Finch-Savage WE. 2013. Temperature, light and nitrate sensing coordinate *Arabidopsis* seed dormancy cycling, resulting in winter and summer annual phenotypes. The Plant Journal 74: 1003–1015.

Forrest J, Miller-Rushing AJ. 2010. Toward a synthetic understanding of the role of phenology in ecology and evolution. Philosophical Transactions of the Royal Society B 365: 3101–3112.

Garrison A. 2022. Adaptation to agriculture in a serious crop weed, weedy radish (Raphanus raphanistrum). PhD Thesis, Michigan State University, USA.

Garrison AJ, Norwood LA, Conner JK. 2024. Plasticity-mediated persistence and subsequent local adaptation in a global agricultural weed (R Spigler and J Wolf, Eds.). Evolution 78: 1804–1817.

Gupta R, Chakrabarty S. 2013. Gibberellic acid in plant: still a mystery unresolved. Plant Signaling & Behavior 8: e25504.

Hall MC, Willis JH. 2006. Divergent selection on flowering time contributes to local adaptation in *Mimulus guttatus* populations. Evolution 60: 2466–2477.

Huberman M. 1941. Why phenology? Journal of Forestry 39: 1007–1013.

Kanomanyanga J, Cussans J, Moss S, Ober E, Liu C, Coutts S. 2025. Adaptation of grassweeds to spring cropping through changes in germination, flowering time and fecundity. Scientific Reports 15: 21492.

Karlsson LM, Milberg P. 2007. Comparing after-ripening response and germination requirements of *Conyza canadensis* and *C. bonariensis* (asteraceae) through logistic functions. Weed Research 47: 433–441.

Landers K. 1995. Vernalization responses in narrow-leafed lupin (*Lupinus angustifolius*) genotypes. Australian Journal of Agricultural Research 46: 1011–1025.

Lang A, Reinhard E. 1961. Gibberellins and flower formation In: Gould R, ed. Advances in Chemistry. *Gibberellins*. ACS Publications, 71–79.

Lenth RV. 2024. emmeans: estimated marginal means, aka least-squares means.

Lu JJ, Tan DY, Baskin JM, Baskin CC. 2014. Germination season and watering regime, but not seed morph, affect life history traits in a cold desert diaspore-heteromorphic annual. PLoS One 9: e102018.

Lu JJ, Tan DY, Baskin CC, Baskin JM. 2016. Effects of germination season on life history traits and on transgenerational plasticity in seed dormancy in a cold desert annual. Scientific Reports 6: 25076.

Main CL, Steckel LE, Hayes RM, Mueller TC. 2006. Biotic and abiotic factors influence horseweed emergence. Weed Science 54: 1101–1105.

Marks M, Prince S. 1981. Influence of germination date on survival and fecundity in wild lettuce *Lactuca serriola*. Oikos: 326–330.

Mennan H, Ngouajio M. 2006. Seasonal cycles in germination and seedling emergence of summer and winter populations of catchweed bedstraw (*Galium aparine*) and wild mustard (*Brassica kaber*). Weed Science 54: 114–120.

Meyer SE, Nelson DL, Carlson SL. 2004. Ecological genetics of vernalization response in *Bromus tectorum* L. (Poaceae). Annals of Botany 93: 653–663.

Michaels SD, Amasino RM. 2000. Memories of winter: vernalization and the competence to flower. Plant, Cell & Environment 23: 1145–1153.

Moon J, Suh S, Lee H, et al. 2003. The *SOC1* MADS-box gene integrates vernalization and gibberellin signals for flowering in *Arabidopsis*. Plant Journal 35: 613–623.

Mutasa-Göttgens ES, Qi A, Zhang W, et al. 2010. Bolting and flowering control in sugar beet: relationships and effects of gibberellin, the bolting gene B and vernalization. AoB PLANTS 2010.

Nakagawa S, Schielzeth H. 2013. A general and simple method for obtaining R2 from generalized linear mixed-effects models. Methods in Ecology and Evolution 4: 133–142.

R Core Team. 2024. R: A Language and Environment for Statistical Computing.

Regehr D, Bazzaz F. 1979. The population dynamics of *Erigeron canadensis*, a successional winter annual. The Journal of Ecology: 923–933.

Rezaee B, Ghasemnezhad A, Zeinali E. 2023. Phenology of evening primrose (oenothera biennis L.) as affected by vernalization and gibberellic acid (GA3). Iranian Journal of Plant Physiology 13: 4589–4597.

Sans F, Masalles R. 1994. Life-history variation in the annual arable weed *Diplotaxis erucoides* (Cruciferae). Canadian Journal of Botany 72: 10–19.

Schramski JA, Sprague CL, Patterson EL. 2021. Environmental cues affecting horseweed (*Conyza canadensis*) growth types and their sensitivity to glyphosate. Weed Science 69: 412–421.

Shah SH, Islam S, Mohammad F, Siddiqui MH. 2023. Gibberellic acid: a versatile regulator of plant growth, development and stress responses. Journal of Plant Growth Regulation 42: 7352–7373.

Smisek A. 1995. Resistance to paraquat in Erigeron canadensis L. MS Thesis, University of Western Ontario, Canada.

Steinmaus SJ, Prather TS, Holt JS. 2000. Estimation of base temperatures for nine weed species. Journal of Experimental Botany 51: 275–286.

Tozzi E, Beckie H, Weiss R, et al. 2014. Seed germination response to temperature for a range of international populations of *Conyza canadensis*. Weed Research 54: 178–185.

Uller T. 2008. Developmental plasticity and the evolution of parental effects. Trends in Ecology & Evolution 23: 432–438.

Van Wychen L. 2022. 2022 Survey of the most common and troublesome weeds in broadleaf crops, fruits & vegetables in the United States and Canada. Weed Science Society of America National Weed Survey Dataset.

Van Wychen L. 2023. 2023 Survey of the Most Common and Troublesome Weeds in Grass Crops, Pasture & Turf in the United States and Canada. Weed Science Society of America National Weed Survey Dataset.

Weaver SE. 2001. The biology of Canadian weeds. 115. Conyza canadensis. Canadian Journal of Plant Science 81: 867–875.

Wickham H. 2016. ggplot2: elegant graphics for data analysis. New York: Springer-Verlag.

Wilson JE. 1981. Effects of formulation and method of applying gibberellic acid on flower promotion in cocoyam. Experimental Agriculture 17: 317–322.

Zeevaart JA. 1983. Gibberellins and flowering. The Biochemistry and Physiology of Gibberellins 2: 333–373.

Zhao L, Li S, Yu Q, et al. 2023. Vernalization promotes GA-mediated bolting initiation via the inhibition of ABA and JA biosynthesis. Agronomy 13: 1251.

