## Supplementary Figures and Table for "Seed vernalization and gibberellic acid interact to affect life cycle type in facultative winter annual Canadian horseweed (*Erigeron canadensis*)"

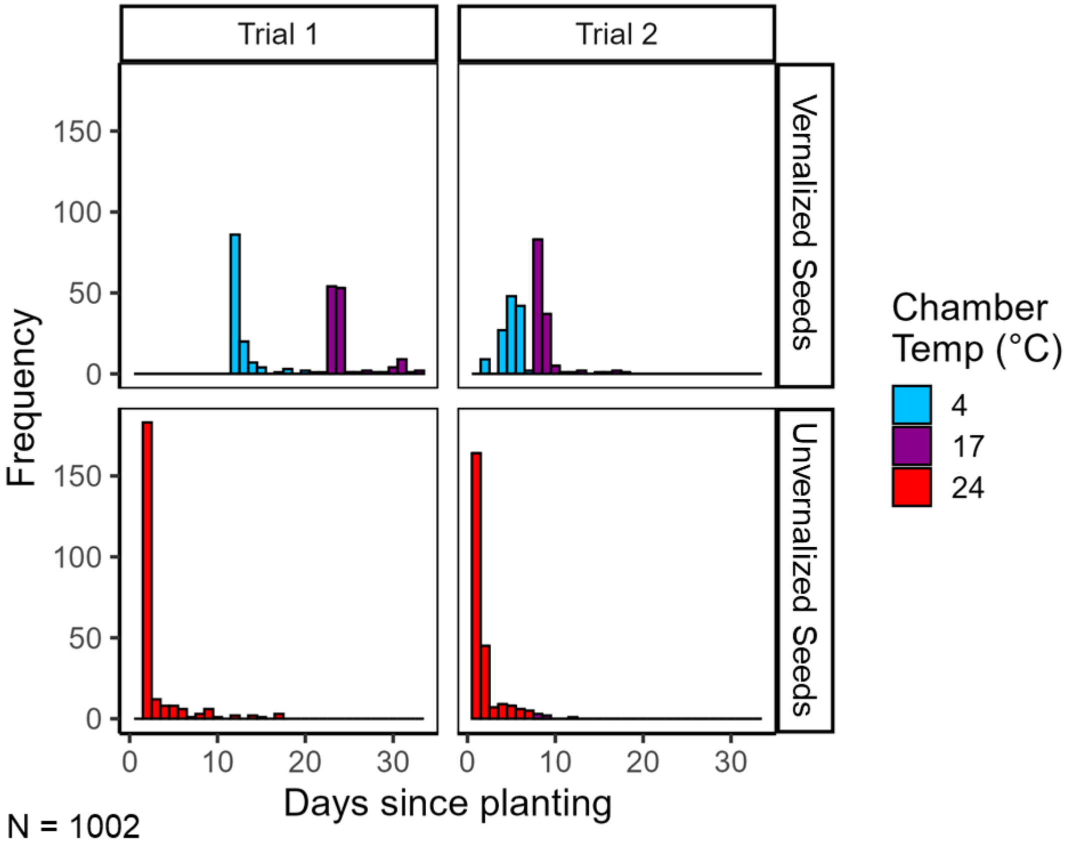


**Figure S1.** Emergence timing by Trial and Vernalization Treatment in the 2023 Growth Chamber Experiment. Bar colors show the temperature of the chamber on the indicated day since planting.

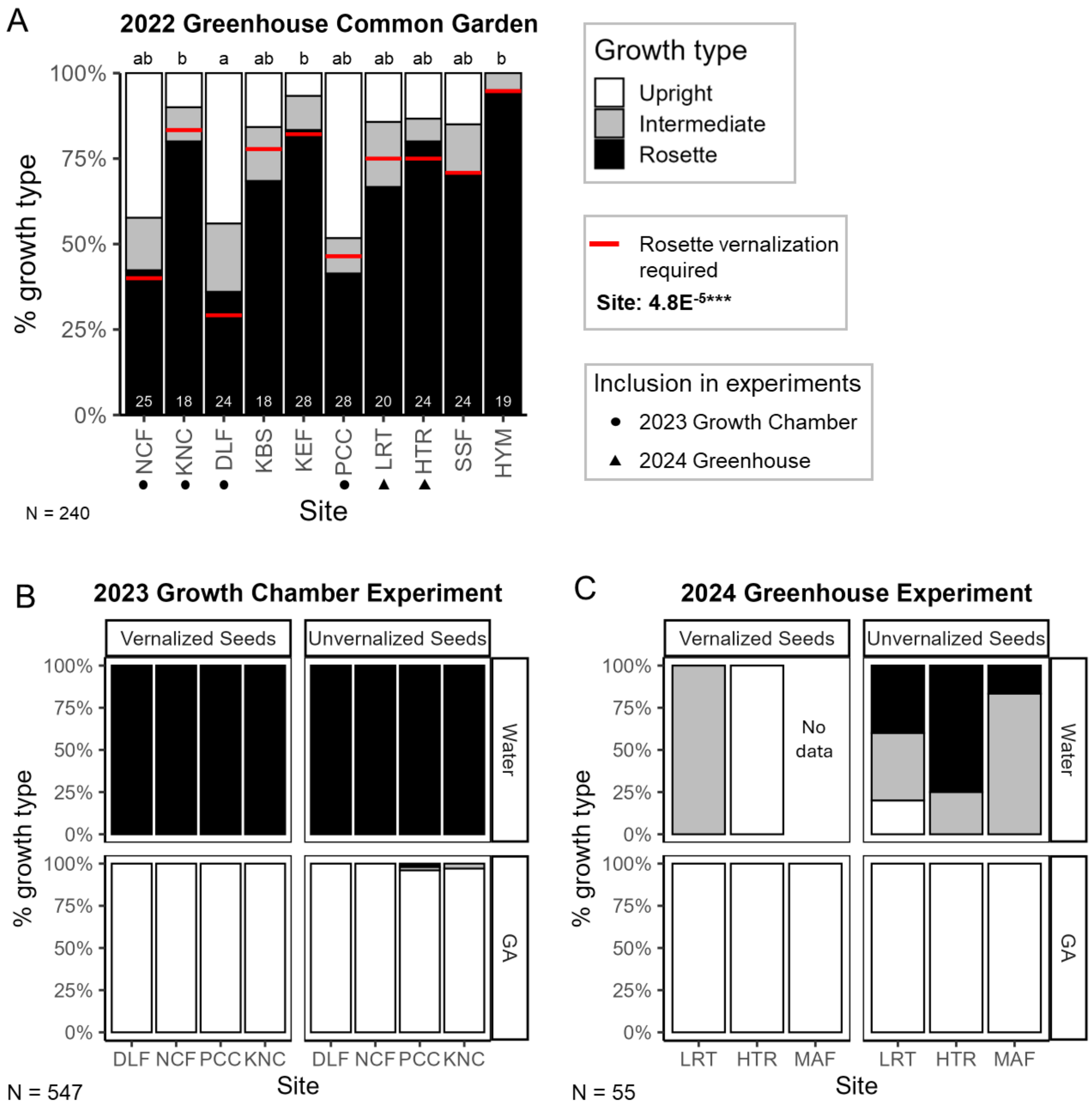


**Figure S2.** Growth type variation by site in greenhouse common garden (A), growth chamber experiment (B), and greenhouse experiment (C). Bars show proportion of plants assigned to upright (white), intermediate (gray), and rosette (black) growth types. In A, sites are sorted first by latitude within 0.1° then by longitude, sample sizes for each site are shown in white text at the bottom of bars, red horizontal lines indicate the proportion of plants that were vernalized after failing to bolt for 11 weeks, sites included in the 2023 Growth Chamber Experiment are indicated by a dot below the site name, and sites included in the 2024 Greenhouse Experiment are indicated by a triangle below the site name. The $P$-value for the effect of Site on vernalization proportion from a Type III Analysis of Deviance is shown and sites that share the same letter are not significantly different in vernalization proportion at $P < 0.05$ from Tukey post-hoc comparisons. In B and C, no site significantly differed from any other site in mean proportion upright type ($P > 0.20$).

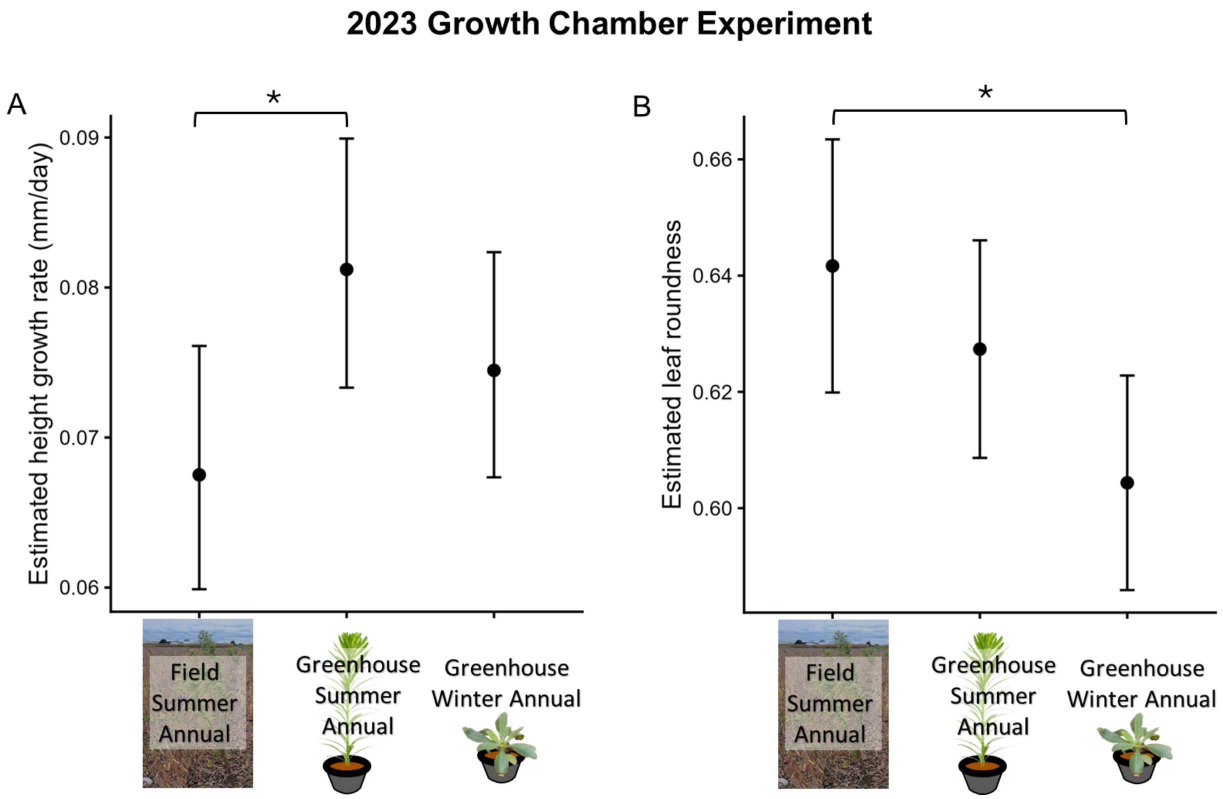


**Figure S3.** Effects of parent plant source type on height growth rate and leaf roundness in the 2023 Growth Chamber Experiment. Points are estimated marginal means after accounting for the other effects in the model and error bars are 95% confidence intervals.

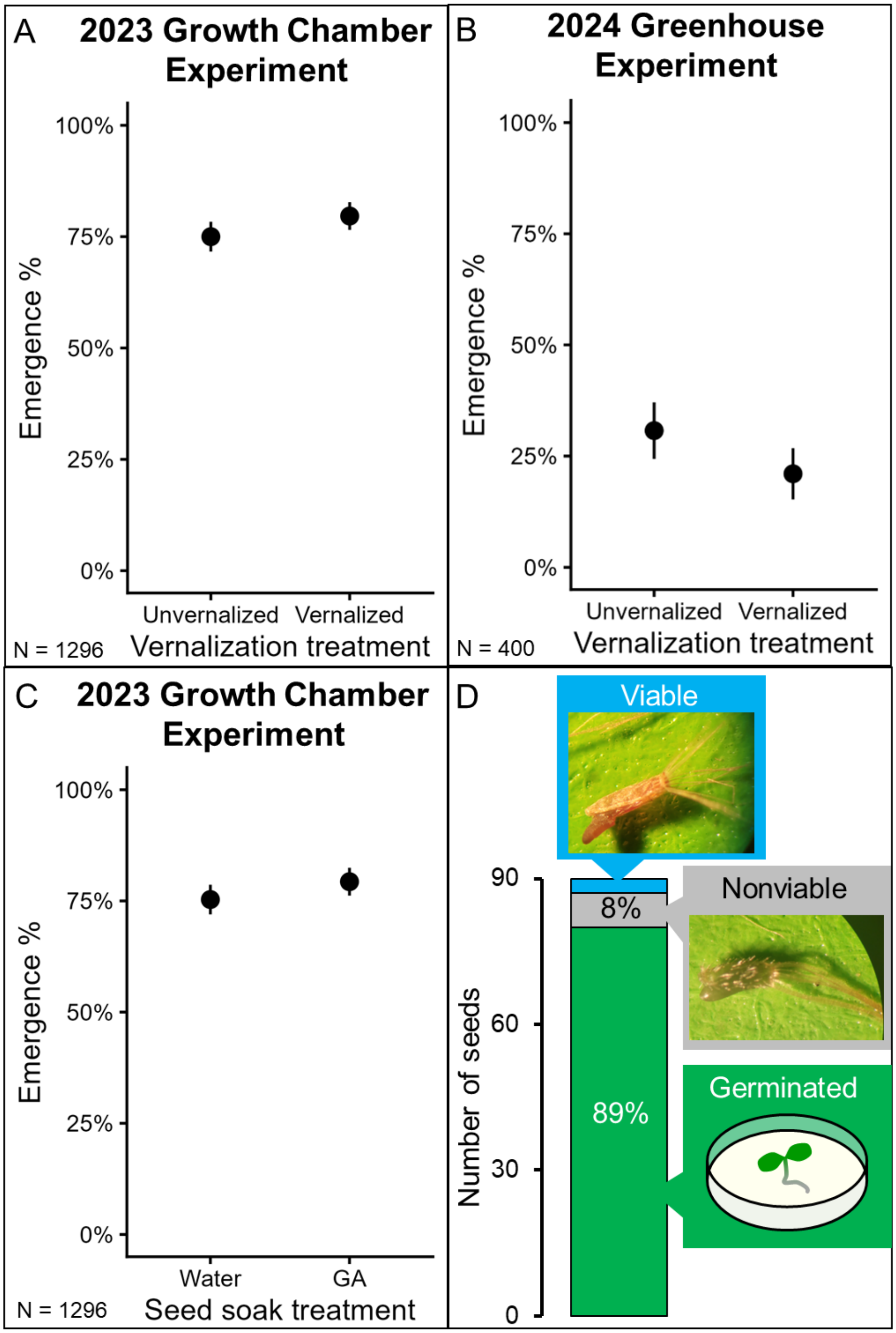


**Figure S4.** Variation in emergence or germination rates. Effects of seed vernalization treatment on emergence rates in 2023 Growth Chamber Experiment (A) and 2024 Greenhouse Experiment (B). Effects of GA seed soak on emergence rate in 2023 Growth Chamber Experiment (C). Initial dormancy state of greenhouse-grown seeds via tetrazolium test (D).

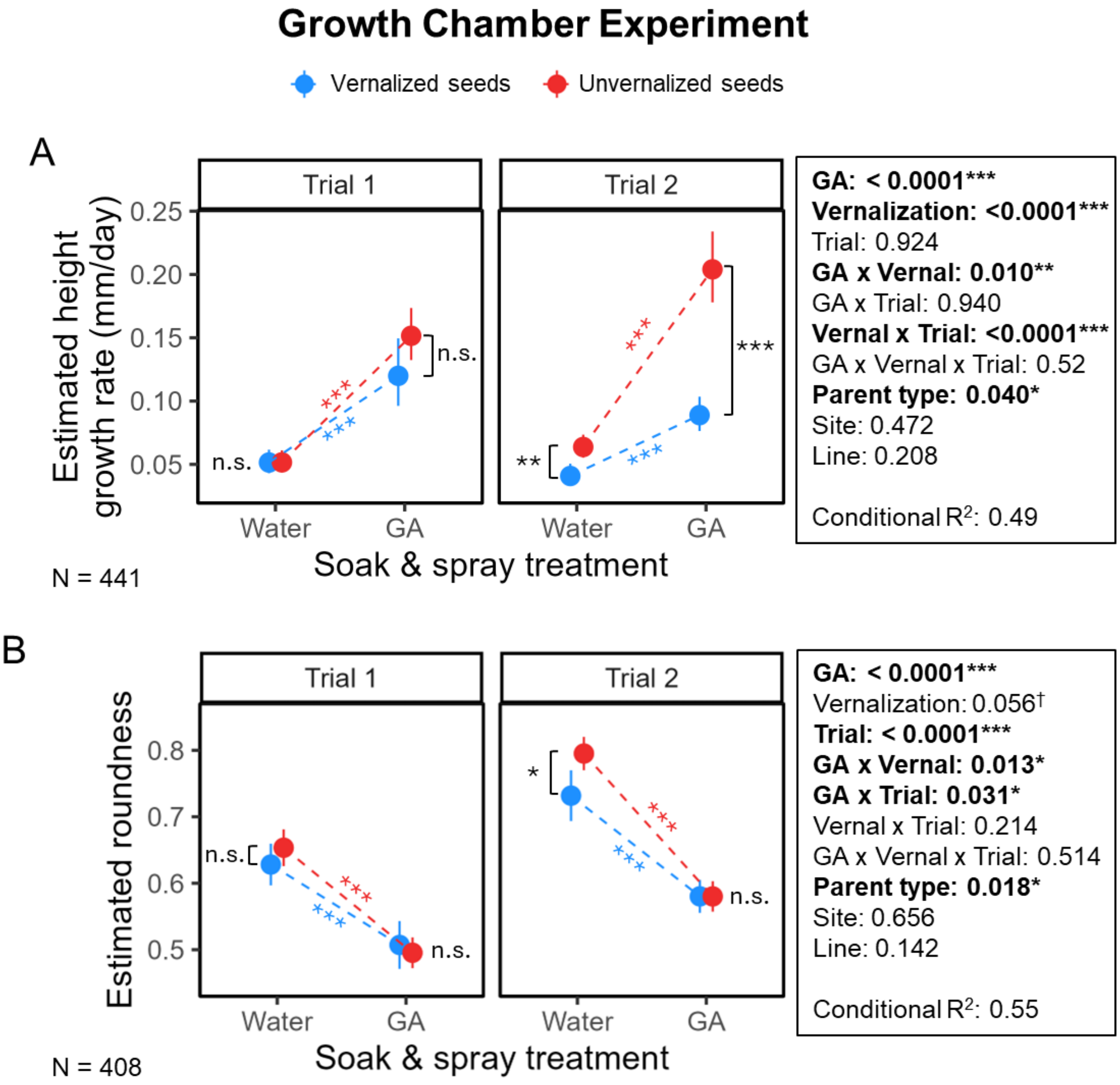


**Figure S5.** Height growth rate and leaf roundness in Growth Chamber Experiment after excluding plants that emerged while in the cold (-4º C). All other details as in Fig. 3.

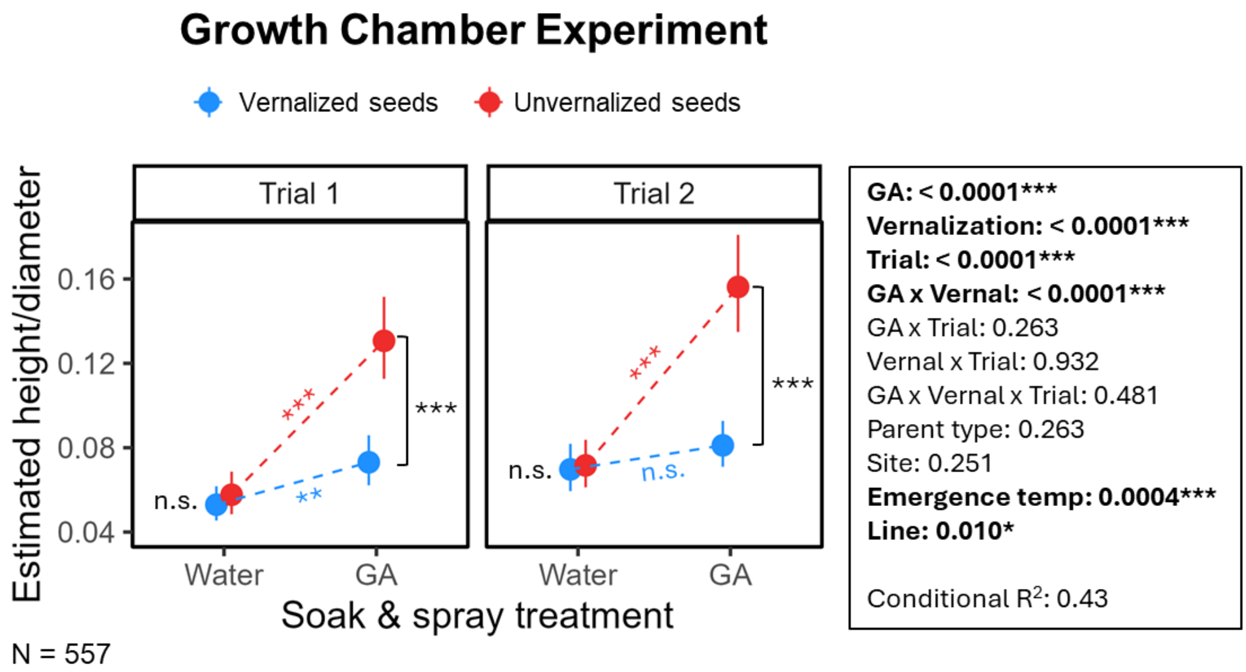


**Figure S6.** Height growth rate measured as plant height per plant diameter in Growth Chamber Experiment. All other details as in Fig. 3.

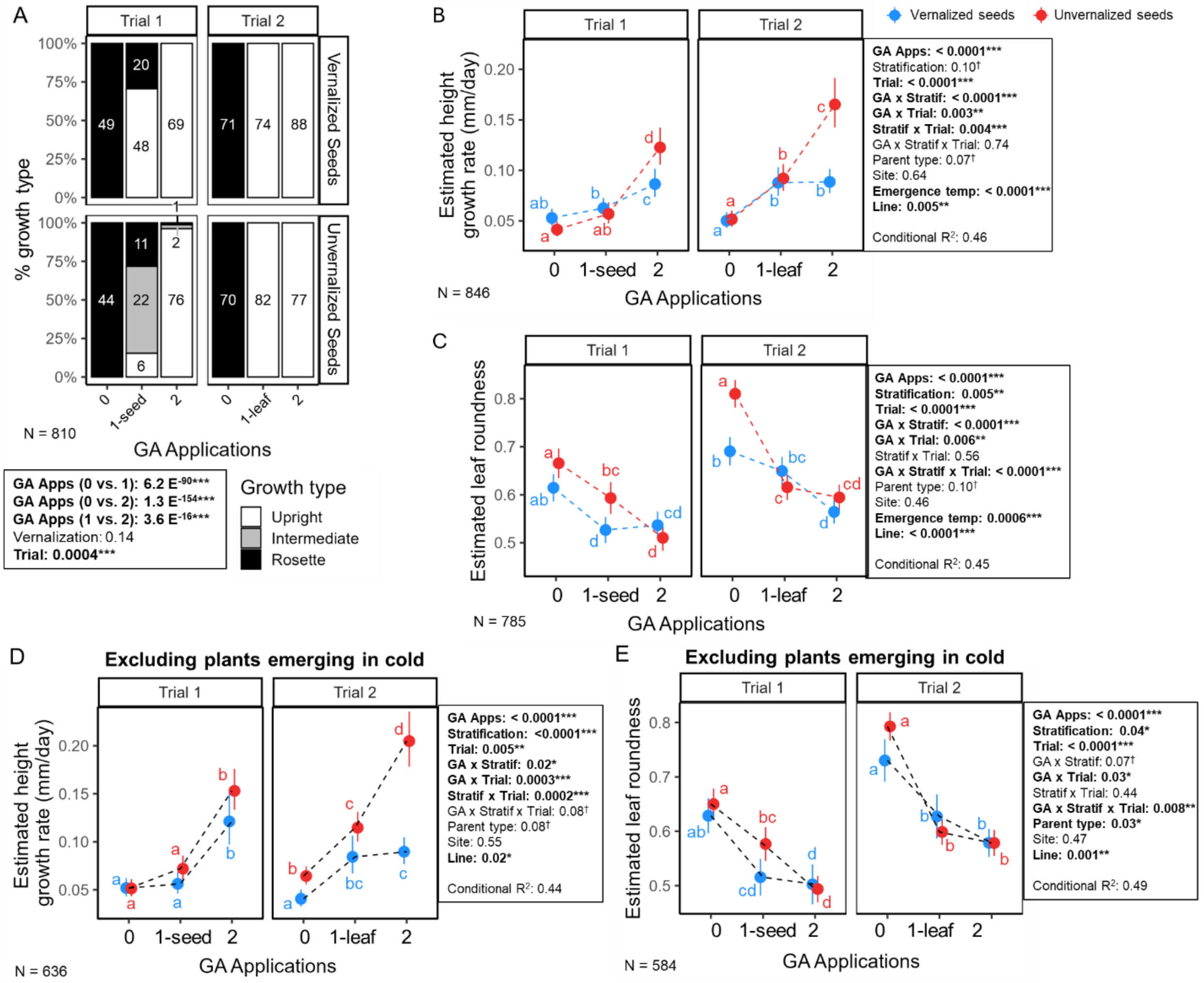


**Figure S7**. Qualitative and quantitative measures of life cycle type in Growth Chamber Experiment showing the single GA application treatments of either only seed soak (Trial 1) or only leaf spray (Trial 2). Growth type assignment (A), height growth rate (B and D), leaf roundness (C and E). Plants that emerged while in the cold (-4ºC) are excluded in D and E. Colors separate vernalization treatments and the slope of dotted lines is the effect of adding GA within vernalization treatment. Panels separate trials. Points are estimated marginal means after accounting for the other effects in the model and error bars are 95% confidence intervals. In A, the box shows Fisher's Exact Test *P*-values for the comparison groups indicated. The boxes in B to E give the *P*-values for all model effects from Type III Analyses of Deviance for fixed effects and likelihood ratio test for the random effect of Line. In B to E, points that share the same letter are not significantly different at $P < 0.05$ from Tukey post-hoc comparisons within panels.

### Supplementary Tables

**Table S1.** Descriptions of all seed source sites used in the study. Ordered by increasing distance from Kellogg Biological Station. Winter average temperatures are from NOAA 30-year U.S. Climate Normals (1981-2010) for closest weather station to field site.

| Site ID | Site Name | Location (Lat, Lon) | Description | Winter avg. temperature (ºC) |
|---|---|---|---|---|
| KBS | Kellogg Bird Sanctuary | Hickory Corners, MI (42.39, -85.38) | Along sunny trail edges in a conservation center | -2.3 |
| KEF | Kellogg Experimental Forest | Augusta, MI (42.35, -85.34) | Along sunny trail edges in a wooded trail system | -2.6 |
| KNC | Kalamazoo Nature Center | Kalamazoo, MI (42.36, -85.58) | Along sunny trail edges near a nature center | -2.6 |
| DLF | DeLano Farms | Kalamazoo, MI (42.36, -85.60) | Within crop rows on an organic mixed vegetable farm | -2.6 |
| PCC | Pierce Cedar Creek | Hastings, MI (42.53, -85.30) | Along sunny trail edges in a mixed prairie and wooded trail system | -3.5 |
| NCF | Natural Cycles Farm | Allegan, MI (42.44, -85.84) | Within crop rows on an organic mixed vegetable farm | -2.7 |
| LRT | Lansing River Trail | Lansing, MI (42.73,-84.51) | Along sunny trail edges in a mixed urban and wooded trail system | -2.7 |
| HTR | Horticulture Teaching & Research Center | Holt, MI (42.67, -84.49) | Within crop rows on an organic mixed vegetable farm | -2.7 |
| MAF | MSU Agronomy Farm | Lansing, MI (42.68, -84.49) | Within crop rows on a conventional soy/corn farm | -2.7 |
| HYM | Hickory Meadows | Traverse City, MI (44.77, -85.66) | Along sunny trail edges and sidewalks at a park | -4.7 |
| SSF | Second Spring Farm | Cedar, MI (44.79, -85.71) | Within crop rows on an organic mixed vegetable farm | -4.7 |
